# TEMPTED: time-informed dimensionality reduction for longitudinal microbiome studies

**DOI:** 10.1101/2023.07.26.550749

**Authors:** Pixu Shi, Cameron Martino, Rungang Han, Stefan Janssen, Gregory Buck, Myrna Serrano, Kouros Owzar, Rob Knight, Liat Shenhav, Anru R. Zhang

**Affiliations:** Department of Biostatistics & Bioinformatics, Duke University, Durham, NC, USA; Duke Microbiome Center, Duke University, Durham, NC, USA; Department of Pediatrics, University of California San Diego, La Jolla, CA, USA; Center for Microbiome Innovation, University of California San Diego, La Jolla, CA, USA; Bioinformatics and Systems Biology Program, University of California San Diego, La Jolla, CA, USA; Department of Statistical Science, Duke University, Durham, NC, USA; Algorithmic Bioinformatics, Department of Biology and Chemistry, Justus Liebig University Giessen, Giessen, Germany; Center for Microbiome Engineering and Data Analysis, Virginia Commonwealth University, Richmond, VA, USA; Department of Microbiology and Immunology, Virginia Commonwealth University, Richmond, VA, USA; Department of Bioengineering, University of California San Diego, La Jolla, CA, USA; Department of Computer Science and Engineering, University of California San Diego, La Jolla, CA, USA; Halicioğlu Data Science Institute, University of California San Diego, La Jolla, CA, USA; Institute for Systems Genetics, New York Grossman School of Medicine, New York University, New York, NY, USA; Department of Microbiology, New York Grossman School of Medicine, New York University, New York, NY, USA; Department of Computer Science, New York University, New York, NY, USA; Department of Computer Science, Duke University, Durham, NC, USA

**Author notes:** P. Shi and C. Martino contributed equally to this work. L. Shenhav and A. R. Zhang contributed equally to this work. Contributing authors.

## Abstract

Longitudinal studies are crucial for understanding complex microbiome dynamics and their link to health. We introduce TEMPoral TEnsor Decomposition (TEMPTED), a time-informed dimensionality reduction method for high-dimensional longitudinal data that treats time as a continuous variable, effectively characterizing temporal information and handling varying temporal sampling. TEMPTED captures key microbial dynamics, facilitates beta-diversity analysis, and enhances reproducibility by transferring learned representations to new data. In simulations, it achieves 90% accuracy in phenotype classification, significantly outperforming existing methods. In real data, TEMPTED identifies vaginal microbial markers linked to term and preterm births, demonstrating robust performance across datasets and sequencing platforms.

## Background

Given the highly dynamic and complex nature of microbial communities, identifying and predicting their time-dependent patterns are crucial to understanding their structure and function. The collection of longitudinal microbiome samples provides a unique opportunity to capture the dynamics of microbial communities and their associations with host phenotypes. However, the nature of longitudinal microbiome data poses several analytical challenges. First, microbiome data are high-dimensional, making dimensionality reduction key in guiding analysis and interpretation. Second, the pattern of intra-host variation may change over time and vary across hosts, making it challenging to extract robust temporal patterns of microbial features [1]. Third, due to inherent practical limitations of longitudinal studies (e.g., missed patient follow-up visits or inconsistent sample collection), multiple hosts often have missing temporal samples, translating into irregular temporal sampling across hosts [2–5].

To investigate longitudinal microbiome data, many studies first use dimensionality reduction methods, such as principal coordinate analysis (PCoA); however, this method analyzes data at the sample level and does not utilize or account for within-subject correlation and temporal structures. In recent years, unsupervised tensor methods have been developed to model longitudinal microbiome data. CTF [1], microTensor [6], TCAM [7], FTSVD [8], and EMBED [9] format temporal microbiome data into tabular tensors and apply tensor decomposition to identify low-dimensional structures. However, these tensor-based methods assume all hosts have the same sampling time points with low-level of or no missingness, which is often unrealistic in clinical settings. Moreover, CTF and microTensor do not account for the continuity in temporal structure, TCAM does not provide dimension reduction for time or samples, while EMBED aims at characterizing temporal structures without offering dimension reduction for hosts or samples. In addition, most of these methods can not transfer the learned low-dimensional representation from training data to independent testing data. Another relevant class of models is the multivariate functional models that can depict feature trajectories. However, they are unsuitable for dimensionality reduction or managing unknown structures in hosts [10]. An alternative to analyzing longitudinal microbiome data is by using supervised methods, which are focused on generative models inferring the dynamics of these communities (e.g., generalized Lotka Volterra) [11–14]. Another example is the mixed-effect type models widely used to quantify intra-host variation, but typically analyze one microbial feature at a time with limited ability to model temporal patterns [15]. While these methods account for the correlation structure induced by repeated measures as well as for sparsity and compositionality, their output does not directly allow the clustering of phenotypes by microbial community dynamics.

Here, we introduce TEMPoral TEnsor Decomposition (TEMPTED), an unsupervised dimensionality reduction tool for high-dimensional temporal data with flexible temporal sampling. TEMPTED formats longitudinal microbiome data into an order-3 temporal tensor with subject, feature, and continuous time as its three dimensions. The tensor is then decomposed into a summation of low-dimensional components, each consisting of a subject loading vector, a feature loading vector, and a temporal loading function (Fig. 1). These loadings provide time-informed dimension reduction and beta-diversity analysis at both the sample and subject levels, and identify corresponding microbial signatures whose temporal trends can aid in discerning host phenotypes. TEMPTED also enables the transfer of the learned low-dimensional representation from training data to unseen testing data, thus facilitating research reproducibility. TEMPTED is unique in that it can handle varying temporal sampling and missing time points, a prevalent issue in longitudinal microbiome studies, without the need of time discretization, sample removal, or sample imputation. Treating time as a continuous variable allows adjacent time points to borrow information from each other, thus reducing the impact of noises and enhancing signals. It is the only dimensionality reduction method currently available for temporal data that offers this flexibility. These unique properties enable TEMPTED to have superior performance in extracting key information in the dataset in data-driven simulations. Further, using TEMPTED we uncover previously undetectable microbial dynamics separating mice with leukemia from healthy ones and pregnancies ending in preterm and term birth.

**Fig 1.**
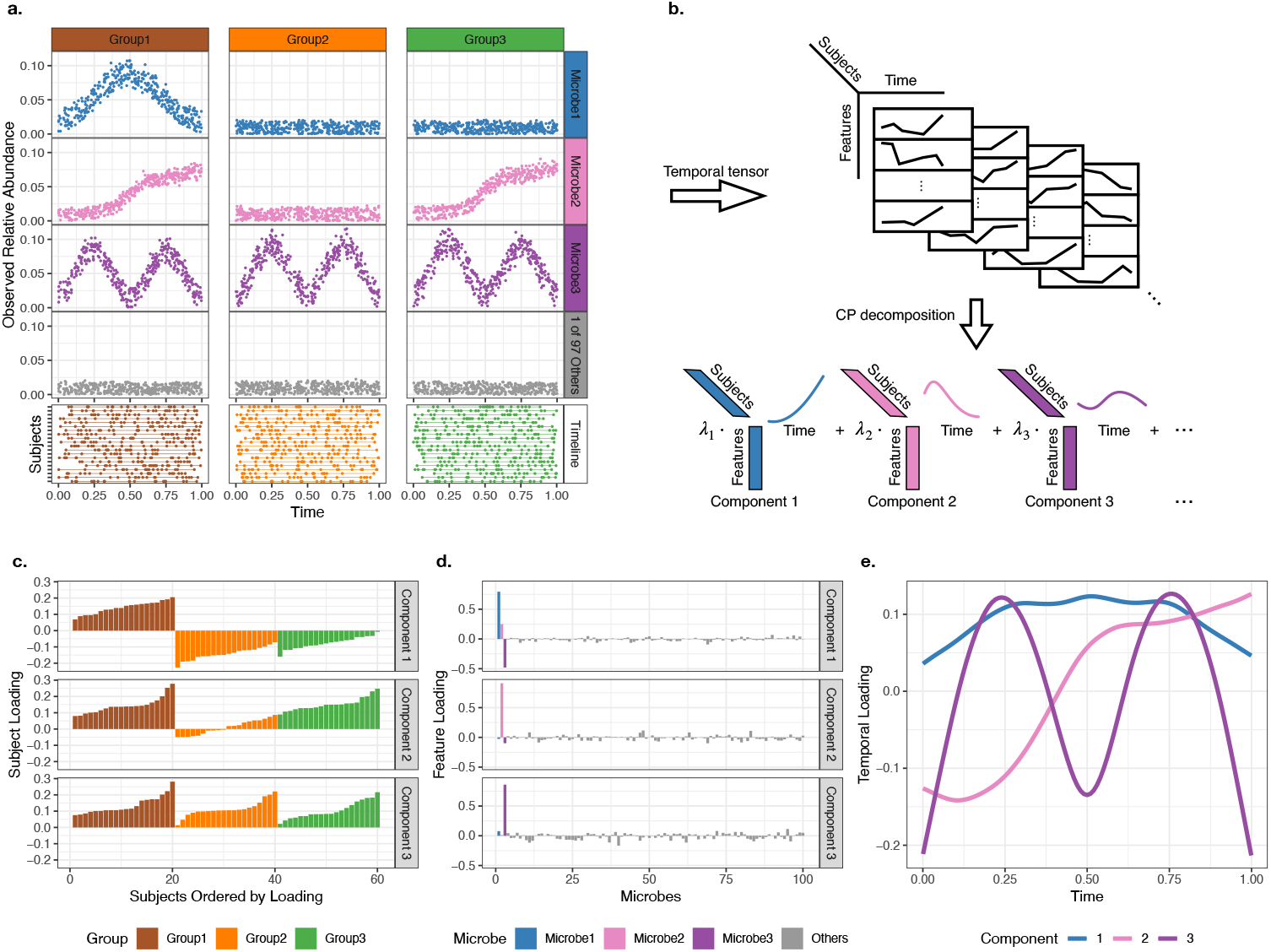
Overview of the TEMPTED algorithm for analyzing multi-subject multi-feature temporal microbiome data. (a) Simulated microbiome count data (see Additional file 1) is transformed into relative abundance and plotted for four representative microbes and three groups of hosts. Microbe 1 has a unique temporal pattern for Group 1, Microbe 2 has a temporal pattern shared by Groups 2 and 3, Microbe 3’s temporal pattern is shared by all groups, and Microbes 4-100 have no temporal patterns. Sampling time points are uniformly distributed and used as is without binning. (b) The observed multi-subject multi-feature temporal data are formatted into a temporal tensor with three modes representing subject, feature (microbe), and time, respectively. TEMPTED reduces the dimension of the temporal tensor by decomposing it into a small number of components, each containing a subject loading vector, a feature loading vector, and a temporal loading function. (c-e) TEMPTED loadings of the first three components from the simulated data. The first component captures the bell-shaped trend of Microbe 1 and separates Group 1 from Groups 2 and 3. The second component captures the increasing trend of Microbe 2 and separates Group 2 from Groups 1 and 3. The third component captures the m-shaped trend of Microbe 3 and does not separate any groups. Microbes 4-100 have low feature loadings in all components.

## Results

### An overview of TEMPTED

Let *i* = 1, …, *n* denote subjects, *j* = 1, …, *p* denote features, and 𝒴_*ijt*_ denote the value of feature *j* from subject *i* at time point 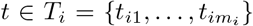. Here, *T*_*i*_ is a subset of interval *T* containing *m*_*i*_ time points and can be different across subjects to accommodate varying temporal sampling and missing time points. TEMPTED allows users to choose their own preferred data normalization and transformation to obtain 𝒴_*ijt*_. As illustrated in Fig. 1b, we adopt the model setting proposed in [8], which decomposes the temporal tensor formed by 𝒴_*ijt*_ using an approximately CANDECOMP/PARAFAC (CP) low-rank structure:

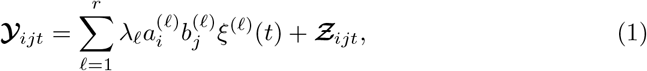

where *r* is the number of low-rank components to approximate the data tensor 𝒴, *λ*_*𝓁*_ quantifies the contribution of each component, 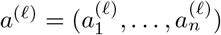 are subject loadings, 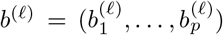 are feature loadings, *ξ*^(𝓁)^(*t*) is the temporal loading that captures the shared temporal patterns among subjects and features, and ***Ƶ***_*ijt*_ includes unexplained remainder terms and measurement errors. Different from FTSVD [8], where *T*_*i*_ is assumed to be identical across all subjects, TEMPTED can accommodate the more common scenario of varying temporal sampling *T*_*i*_ across subjects. Our objective is to estimate *λ, a*^(𝓁)^, *b*^(𝓁)^, and *ξ*^(𝓁)^(*t*) while requiring *ξ*^(𝓁)^(*t*) to be smooth. For details of the assumptions on *ξ*(*t*) and the algorithm for estimation, see Methods. The subject loadings *a*^(𝓁)^ can be used for subject-level beta analysis such as classifier training. The feature loading *b*^(𝓁)^ quantifies the contribution of Feature *j* to Component 𝓁. They can be used as the weights vector to aggregate all *p* features into the following subject-specific trajectory corresponding to Component 𝓁:

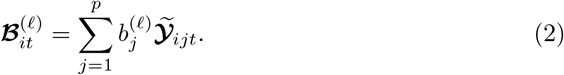

Here, the vector 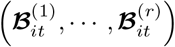 also serves as an *r*-dimensional representation of Sample *t* from Subject *i*, which can be used to construct Euclidean distance and perform sample-level beta analysis. The definition of (2) is generic and can be applied to any longitudinal temporal data.

Since most microbiome data from 16S or shotgun metagenomic sequencing are compositional, researchers are often interested in the relative abundance of one group of microbes versus another. In this scenario, the users can zoom into a small number of features most relevant to each component by specifying a quantile cutoff for the feature loadings, and construct trajectories of log-ratio abundance of top over bottom ranking features:

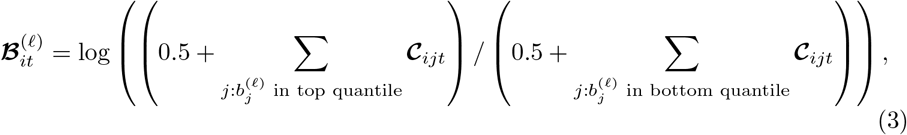

where **𝒞**_*ijt*_ is the read count of Feature *j* in the *t*th sample of Subject *i*. Details of the pseudocount chosen in this log ratio transformation can be found in Methods.

### Data-driven simulation

We evaluated TEMPTED’s ability to perform phenotype discrimination using two data-driven simulations. The first simulation is based on the ECAM dataset [2], which sampled the gut microbiome of infants delivered vaginally versus by C-section during the first two years of life (Additional file 1: Fig. S6). This dataset was chosen due to previously observed differences in longitudinal trajectories between delivery modes [1, 2]. The second simulation utilizes the FARMM [16] dataset that comprises daily fecal microbiome samples collected over 15 days from 30 individuals equally divided into three dietary categories—vegan, omnivore, and exclusive enteral nutrition (EEN) without dietary fiber—with all subjects receiving antibiotic treatment during days 6 to 8 (Additional file 1: Fig. S7). This dataset was chosen due to its use of metagenomics sequencing, and previously observed differences between EEN diet and the other two in the recovery of microbiome after antibiotic treatment [6, 16]. We evaluated TEMPTED and alternative computational methods on their ability to differentiate host phenotypes based on microbial dynamics at the subject level through precision-recall (PR) of classification and the sample level through PERMANOVA F-statistic (see Methods).

These evaluations were performed across random subsets of samples in order to simulate sparse and varying temporal sampling to assess the impact of different sampling densities. First, at the subject level, we evaluated the performance of TEMPTED as compared to CTF, microTensor, FTSVD, and TCAM, since methods like PCoA, including Bray-Curtis, Unifrac, and Weighted Unifrac are limited to sample-level analysis. TEMPTED outperforms all methods and reduces the AUC-PR error of host-phenotype classification by more than 50% compared to CTF and microTensor, and this superiority is maintained even when other methods utilize more time points (Fig. 2a,c). Among these methods, TEMPTED and TCAM are the only ones capable of transferring the learned low-dimensional representation from training to testing data (see Methods) and performing out-of-sample prediction, although TCAM cannot handle missingness in time points (Fig. 2a,c). Second, at the sample level, TEMPTED outperforms all existing methods in phenotype differentiation across all sampling densities (Fig. 2b), while TCAM does not provide sample-level dimension reduction. It is also important to note that among all the non-PERMANOVA methods, TEMPTED is the only method treating time as a continuous variable without discretization or imputation. While CTF and microTensor allow missingness in the time points, they require discretizing time into intervals. On the other hand, TCAM and FTSVD require temporal sampling to be identical across subjects without any missing time points. To use these methods for highly varying temporal sampling, as in our ECAM-based simulation, without sample imputation, we transformed time into the order of samples (Fig. 2) or monthly intervals (Additional file 1: Fig. S5)

**Fig 2.**
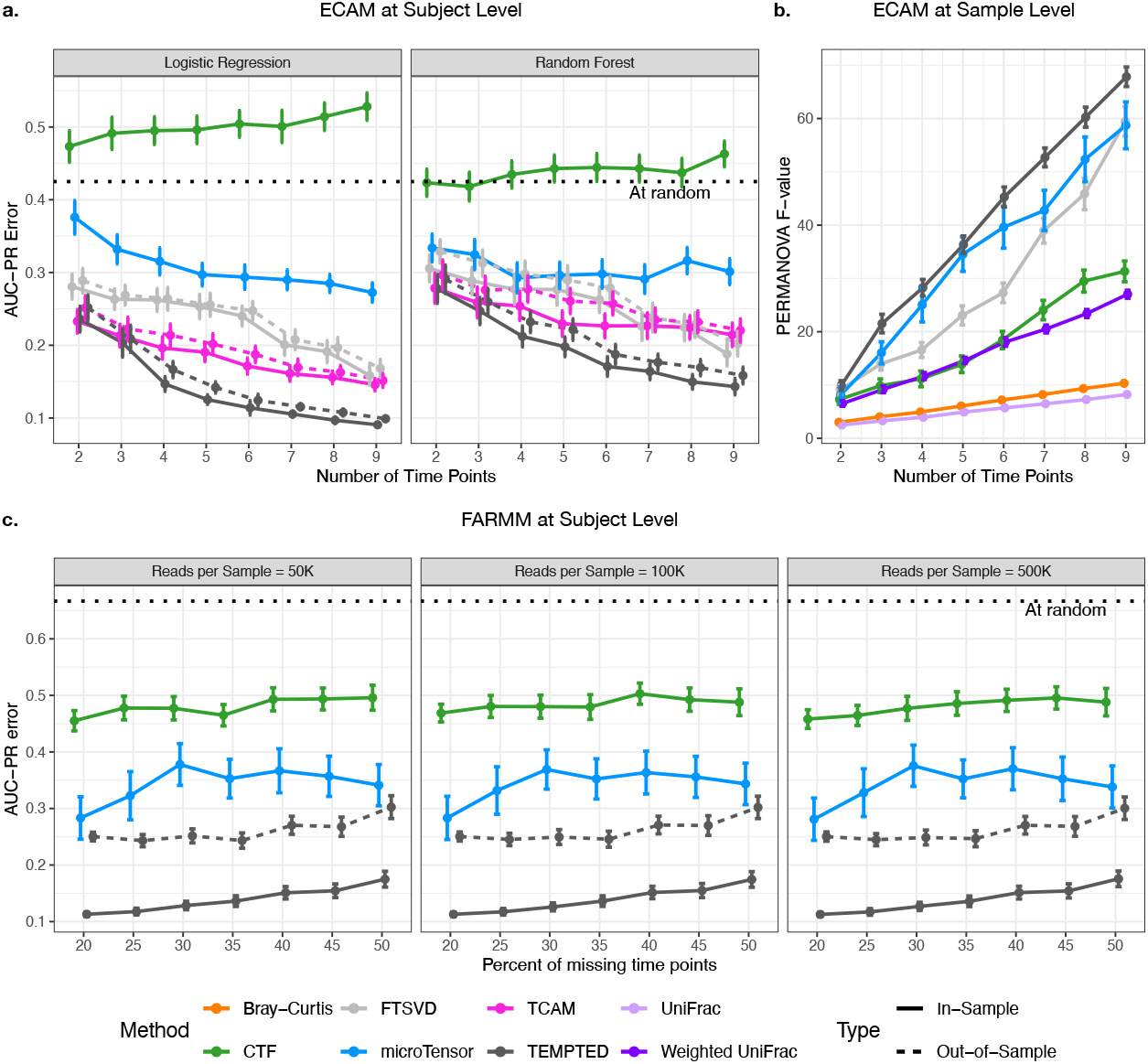
TEMPTED outperforms CTF, microTensor, TCAM, FTSVD, and PCoA in identifying group structures. TEMPTED demonstrates superior performance in reducing host-level phenotype discriminatory error by more than 50% compared to methods capable of handling missing time points (a, c) and improves the sample-level group discriminatory power (b). Sample-level discriminatory power is quantified by PERMANOVA pseudo F-statistic based on the sample-level Euclidean distance constructed using the first two components from each method (b) (for TEMPTED see Eq. (2)). Host-level group classification error is quantified by AUC-PR (1 - area under the precision-recall curve) with the first two components of each method as predictors. Both logistic regression and random forest classifiers were employed and shown in (a), and the results from the better of the two classifiers were shown in (c). Dimension reduction is performed in-sample and out-of-sample (See Methods) respectively, and group labels are predicted using leave-one-out for logistic regression and out-of-bag for random forest. The methods were applied to two datasets: the ECAM infant fecal microbiome data (a, b), which distinguishes between infants delivered vaginally (N-subject=23) and by cesarean section (N-subject=17); and the FARMM dataset (c), which distinguishes between EEN diet (N-subjects=10) and vegetarian or omnivore diet (N-subject=20). Error bars represent 1.96 standard errors. For ECAM-based simulation, we randomly choose a given number of samples from each subject such that CTF, microTensor, FTSVD and TCAM can use the order of the infant age as time variable to form a tensor with no missing values, while TEMPTED uses the infant age as is. For FARMM-based simulation, we randomly drop samples from 15 time points to achieve different percent of missingness, which CTF and microTensor can manage but TCAM and FTSVD cannot. EMBED was not included in the benchmarking because it does not provide host-level or sample-level beta diversity analysis. Different reads per sample are obtained by resampling reads in each sample.

### Case studies

To further examine the performance of TEMPTED on real data, we applied it to three publicly available datasets. First, we used a mouse study of Acute Lymphoblastic Leukemia (ALL), the most common form of childhood cancer with high genetic predisposition [3]. Fecal microbiome samples were longitudinally sampled from wildtype and predisposed (Pax5+/- genotype) mice that were raised in a specific pathogen-free environment and transferred to a conventional facility at early adulthood to resemble children’s transition into kindergarten [3] (Additional file 1: Fig. S8). While both TEMPTED and existing methods could successfully recover mouse genotype from the microbiome (wild type vs. Pax5+/-), existing methods failed to consistently predict the onset of ALL from microbial profiles (Additional file 1: Figs. S11-13, Tables S1-2). In contrast, TEMPTED was the only method that clearly distinguished healthy mice from those that developed ALL while associating the difference to specific ASVs (Fig. 3a-c, Additional File 1: Fig. S14, Additional File 2). Additionally, TEMPTED identified ASVs that clearly separate wildtype and Pax5+/- mice (Fig. 3b-c, Additional file 1: Fig. S15). As there is currently no reliable biomarker for ALL onset, this finding might facilitate lead time for therapy. Furthermore, TEMPTED could correlate disease onset to just 12 out of 1065 ASVs. Several of the ALL onset associated ASVs have been associated previously with leukemia and other cancers in mice, in particular those classified in the Acetatifactor genus [17–19].

**Fig 3.**
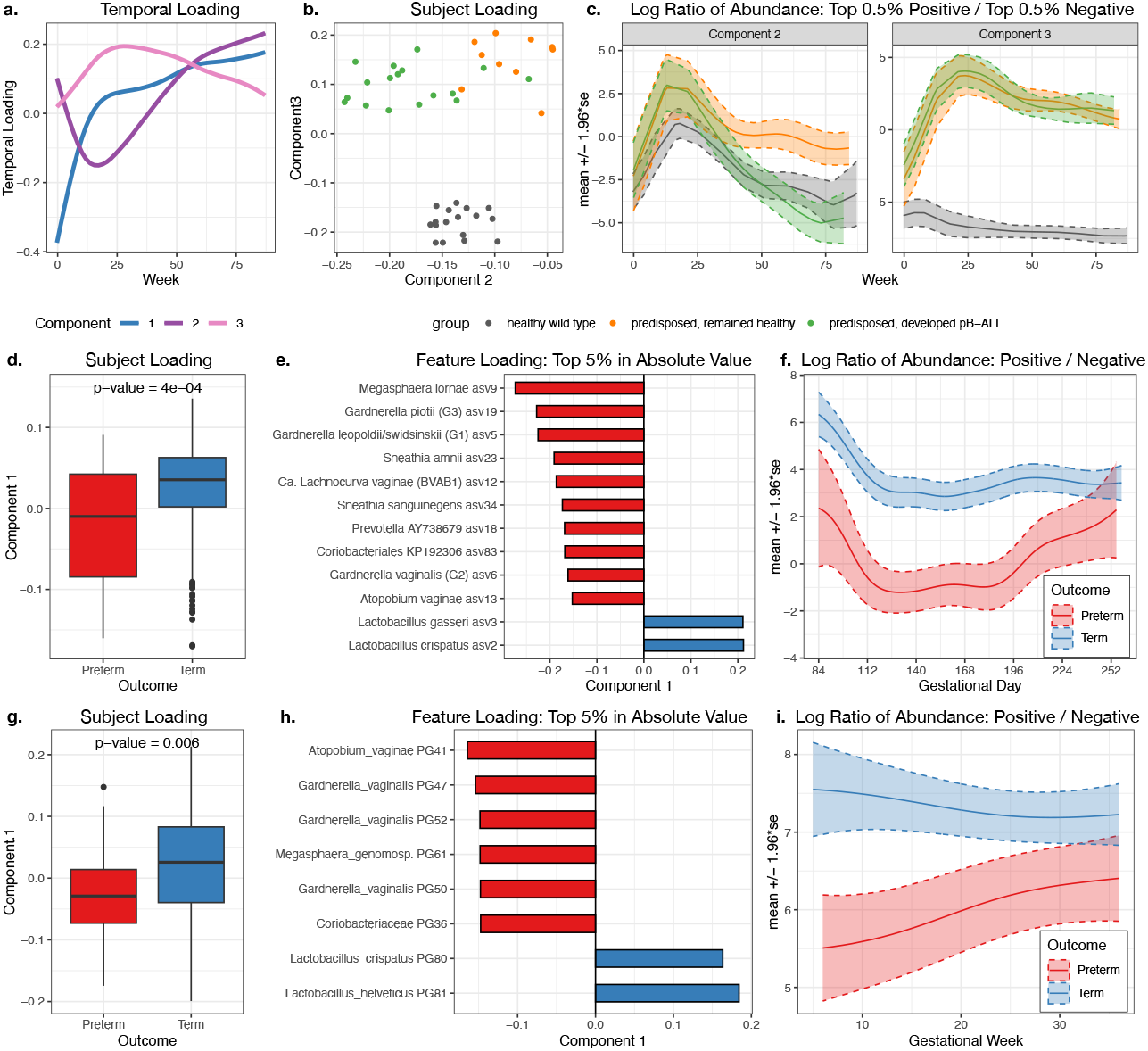
TEMPTED reveals latent trajectories distinguishing health phenotypes. (a)-(c) show the results of applying TEMPTED to a mouse study on leukemia (17 wild-type mice, 27 Pax5+/-mice of which 17 developed leukemia)3. (a) Temporal loadings capture major temporal patterns in the central-log-ratio-transformed microbial abundance. (b) Subject loadings separate genotypes in Component 3 and disease statuses in Component 2. (c) Log ratio of the total abundance of the top 0.5% ASVs over the bottom 0.5% ASVs, where 1065 ASVs are ranked by feature loadings of Components 2 (left) and Component 3 (right), respectively. The log ratio from Component 2 diverges between disease statuses for Pax5+/-mice at around 40 weeks after antibiotic treatment. The log ratio from Component 3 shows distinct temporal patterns between genotypes. (d-f) Results of applying TEMPTED to ASVs in the vaginal microbiome during pregnancy (141 term and 49 preterm) [4, 5]. (g-i) Results of applying TEMPTED to MAGs in the vaginal microbiome during pregnancy (111 term and 35 preterm) [4, 20]. (d,g) Subject loadings separating pregnancies resulting in term and preterm deliveries in Component 1. The P-value is from the Wilcoxon rank sum test. (e,h) Top ASVs and MAGs ranked by the absolute value of feature loadings of Component 1. Features with negative and positive loadings are associated with preterm birth and term birth, respectively. (f,i) Log ratio of the abundance of the ASVs with top positive loadings and top negative loadings in (e,h), separating pregnancies resulting in term and preterm deliveries.

Next, we applied TEMPTED to two independent studies that longitudinally sampled the vaginal microbiome during pregnancy: one study sequenced shotgun metagenomics and used metagenome-assembled genomes (MAGs) [4, 20] and while the second sequenced 16S rRNA genes and used ASVs [4, 5] (Additional file 1: Fig. S9-10). Remarkably, TEMPTED consistently captured differences in the vaginal microbiome trajectories of pregnancies that resulted in term versus preterm birth in both datasets. Notably, this differentiation between term and preterm births is based on the dynamics of 8 leading microbial features in each study with a significant overlap (Fig. 3e, h), which can potentially serve as a biomarker to detect preterm birth early in pregnancy (Fig. 3f, i). Of note, the dynamics of these features are required to differentiate between term and preterm birth as this separation was not found when performing differential abundance analysis using ALDEx2 [21] at any single time point (Additional Files 3-4; Methods). The identified leading ASVs and MAGs both associate Lactobacillus spp. and specifically Lactobacillus crispatus with term birth, and associate Gardnerella, Megasphaera, and Atopobium vaginae with preterm birth (Fig. 3e, h), consistent with existing literature [4, 20, 22]. Overall, our results highlight the robustness of TEMPTED in multiple studies and sequencing technologies in uncovering microbial dynamics underlying host phenotypes.

## Discussion

Despite its many strengths, users of TEMPTED should also be aware of several limitations. Like other unsupervised dimensionality reduction methods such as PCA, PCoA, CTF, microTensor, FTSVD, and TCAM, TEMPTED extracts prominent structures from the data, but does not guarantee the capture of the phenotype of interest in its leading components or a single component or look for differential temporal trends between host groups. TEMPTED’s smoothness assumption (see Methods) on the temporal loading allows it to extract key temporal trends beneath the noisy observed data, but such an assumption also makes it unsuitable for change point detection. Currently, TEMPTED cannot handle individual missing entries in the data tensor, which could be addressed in future work. In addition, TEMPTED does not guarantee its rank-*r* decomposition to have the best reconstruction accuracy because it obtains its low-rank structure for each component sequentially to achieve the uniqueness of the decomposition instead of looking for the best rank-r approximation for a given rank. Nevertheless, TEMPTED has a flexible model setting that accommodates a wide range of high-dimensional temporal data, making it a valuable and powerful tool for research beyond longitudinal microbiome studies.

## Conclusions

TEMPTED was designed to address an important unmet need in the rapidly evolving field of microbiome research – namely unsupervised dimensionality reduction for longitudinal microbiome data that account for the temporal order of samples and accommodates varying temporal sampling schemes and missing values. By leveraging temporal patterns shared across hosts and features, TEMPTED efficiently extracts important low-dimensional structures from high-dimensional longitudinal data, identifies major temporal dynamics and key contributing features, facilitates beta-diversity analysis at both sample and subject levels, and allows the transfer of the learned low-dimensional representation from training data to unseen datasets.

We demonstrated the utility of TEMPTED using data-driven simulations and real data applications. First, using data-driven simulations based on data sequenced by two technologies (ECAM by 16S and FARMM by shotgun metagenomics), we showcase that TEMPTED outperforms existing methods by a large margin in the ability to extract low-dimensional data structures associated with phenotype differences. Second, using three publicly available datasets on different ecosystems (vaginal microbiome and gut microbiome), we showcase that TEMPTED uncovers biological signals that go undetected by existing methods and produce reproducible results across studies and data types (e.g., 16S and metagenomic sequencing). The microbial features identified by TEMPTED are strong indicators of phenotype differentiation and thus can be leveraged in the design of microbiome-based biomarkers. Finally, its flexibility in accommodating varying temporal sampling, and ability to transfer the learned low-dimensional representation from training to testing data, make it a highly practical tool for microbiome research, promoting the research reproducibility. The flexible model setting of TEMPTED also makes it applicable to a wide range of high-dimensional temporal data, potentially benefiting more research beyond longitudinal microbiome studies.

## Methods

In this section, we provide details of our methods, including data preprocessing, the TEMPTED algorithm, tensor reconstruction, and how the dimension reduction learned from training data can be transferred to new testing data. Additionally, we present details of case studies (Pax5 Mice Leukemia data, shotgun metagenomic vaginal data, 16S Vaginal data), and data-driven simulation studies.

### Microbiome Data Preprocessing

Here, we present the data transformation we adopted for microbiome sequencing data specifically to address their compositionality and highly skewed distribution. For other sources of data, users can choose their desired transformation and normalization before using TEMPTED. Let *i* = 1, …, *n* denote subjects, *j* = 1, …, *p* denote features, and **𝒞**_*ijt*_ denote the observed read count of feature *j* from subject *i* at time points 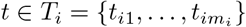 We apply the centered-log-ratio (CLR) transformation to read counts added by .5 [23, 24]

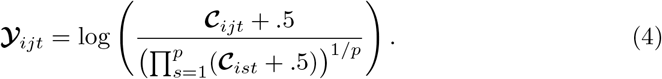

Adding the pseudocount .5 instead of other values is theoretically justified by [24]. [24].showed that log(**𝒞**_*ijt*_ + *α*_*i*_*/*2) has the smallest bias in the estimation of log(mean of **𝒞**_*ijt*_) when (**𝒞**_*i*1*t*_, …, **𝒞**_*ipt*_) follow Dirichlet-Multinomial distribution with *α*_*i*_ being the overdispersion parameter. Since microbiome sequencing count data are generally equally or more dispersed than multinomial distribution [25], when the estimation of *α*_*i*_ is difficult, we opt to estimate *α*_*i*_ with 1, leading to the pseudocount of .5. In Additional file 1: Fig. S1, we also compare the pseudocount of 0.5 with 0.1 and 1. While using 0.1 yields slightly worse performance in subject-level and sample-level dimension reduction, choosing between 0.5 and 1 has little difference. It is also worth noting that among the methods we benchmarked against in our simulation analyses, TCAM, CTF, and FTSVD all require log transformation of the count data. microTensor models count data directly without adding pseudocounts and log transformation, yet it is still outperformed by our method.

### TEMPTED algorithm

The goal of TEMPTED is to obtain estimates of *λ*_𝓁_, *a*^(𝓁)^, *b*^(𝓁)^ and *ξ*^(𝓁)^(*t*) in the approximately CP low-rank structure (1). To overcome the issue of scaling identifiability (e.g., (*λ*_*l*_, *a*^(*l*)^, *b*^(*l*)^, *ξ*^(*l*;)^(*t*)) and (*x*_1_*λ*_*l*_, *x*_2_*a*^(*l*)^, *x*_3_*b*^(*l*)^, *x*_4_*ξ*^(*l*)^(*t*)) essentially represent the same component whenever *x*_1_*x*_2_*x*_3_*x*_4_ = 1), we opt to estimate each component 𝓁 sequentially for 𝓁 = 1, …, *r* by minimizing the following objective function (5).

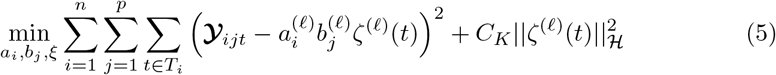

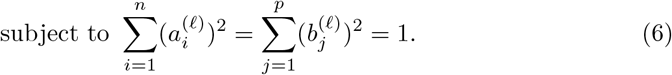

While the sequential estimation of components does not guarantee the smallest reconstruction error, it preserves the uniqueness of each component regardless of the chosen rank *r* and offers the significant advantage of allowing users to explore additional components without impacting those previously obtained.

In (5), 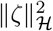 is the *reproducing kernel Hilbert space (RKHS) norm* of function *ζ*(*t*) with the rescaled Bernoulli polynomial as the reproducing kernel,

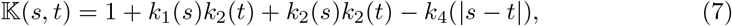

where 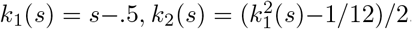, and 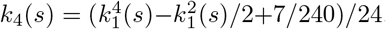. This kernel guarantees *ζ*(*t*)s to be absolutely continuous and squared-integrable in its second-order derivative, and *C*_*K*_ is a tuning parameter controlling the smoothness of *ζ*(*t*)s. Such smoothness assumption does not require the temporal trends themselves to be polynomials. Commonly seen temporal trends such as monotone, unimodal, bimodal, seasonal or circadian trends, can all satisfy this smoothness assumption. It is worth noting that the smoothness assumption is on the underlying structure of the tensor, not the observed tensor itself. Thus, variation of microbiome data due to noises does not necessarily violate our smoothness assumptions.

Due to the substantial differences in abundance among bacterial taxa across most time points, we offer an optional step we referred to as “mean subtraction”, to extract and remove this time-invariant structure from 𝒴 to improve the efficiency of our algorithm when this structure is not of interest. Specifically, we calculate the average of 𝒴_*ijt*_ across time points *t*, resulting in 𝒴_*ij·*_, which forms the matrix 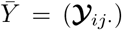Then, we calculate the first singular component of 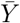 as 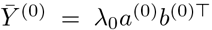, and subtract 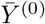 from 𝒴_*t*_ for each *t*. This optional mean subtraction step reduces the effect of highly abundant bacteria taxa and other time-invariant factors that may confound downstream analysis. We adopted this mean subtraction in all our simulation analyses in the main text and real data analyses. In Additional file 1: Section S3, we provide more insights into the effect of mean subtraction through simulation based on the ECAM data and FARMM data. In summary, we found that the reconstruction accuracy of TEMPTED with mean subtraction at rank *r* is comparable to but slightly better than that of TEMPTED without mean subtraction at rank *r* + 1 (Additional file: Fig. S3).The ability of TEMPTED to capture group differences is greatly reduced without mean subtraction, but this reduction can be compensated by adding another component to the classification and is highly dependent on the dataset (Additional file 1: Fig. S4). These results suggest the possibility of TEMPTED capturing the mean structure in some of its components depending on its significance in the dataset. Depending on whether the mean structure contains useful information for downstream analyses, users can decide if mean subtraction is needed before applying TEMPTED.

Set 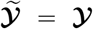 after mean subtraction. For each component 𝓁 = 1, …, *r* sequentially, we perform the following Steps 1 to 3 to estimate *a, b*, and *ζ*(*t*).

Step 1: (Initialization) Initialize 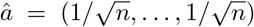 Set 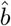 as the first left singular vector of mode-2 matricization of 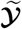 : i.e., the *p*-by- 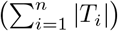 matrix with 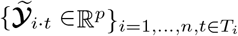 as its columns.

Step 2: (Estimation of loadings) To estimate the loadings, we minimize the following function by iteratively updating 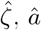 and 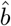 respectively until convergence:

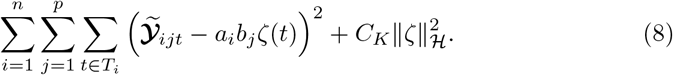

- Update 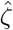 by applying kernel ridge regression to solve

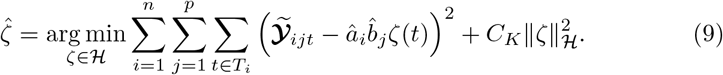 The details of this update are described in the next section.
- Update *â* = (*â*_1_, …, *â*_*n*_) by

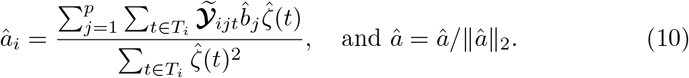
- Update 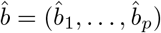 by

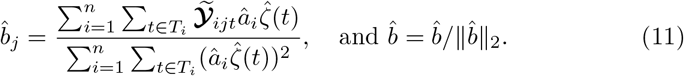

Step 3: (Subtracting previous components) Normalize 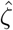 to 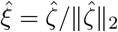. Update 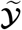 by

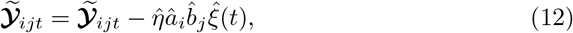

where 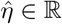 is obtained by solving the following least squares problem:

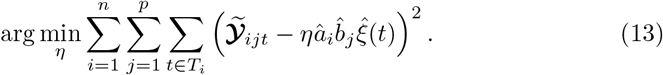

Step 4: After obtaining 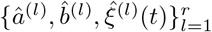 by sequentially running Steps 1-3, we estimate *λ* = (*λ*_1_, …, *λ*_*r*_) is estimated via the following least squares problem:

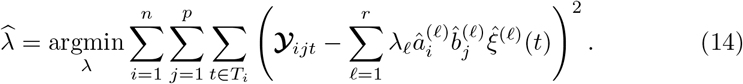

### Kernel Ridge Regression

By the presenter theorem [26], the solution to (9) must be a linear combination of 𝕂(*·, s*) for 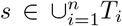 *T*_*i*_, and the weight of this linear combination can be solved by kernel ridge regression. Specifically, we introduce the concatenated observation vector and kernel matrix:

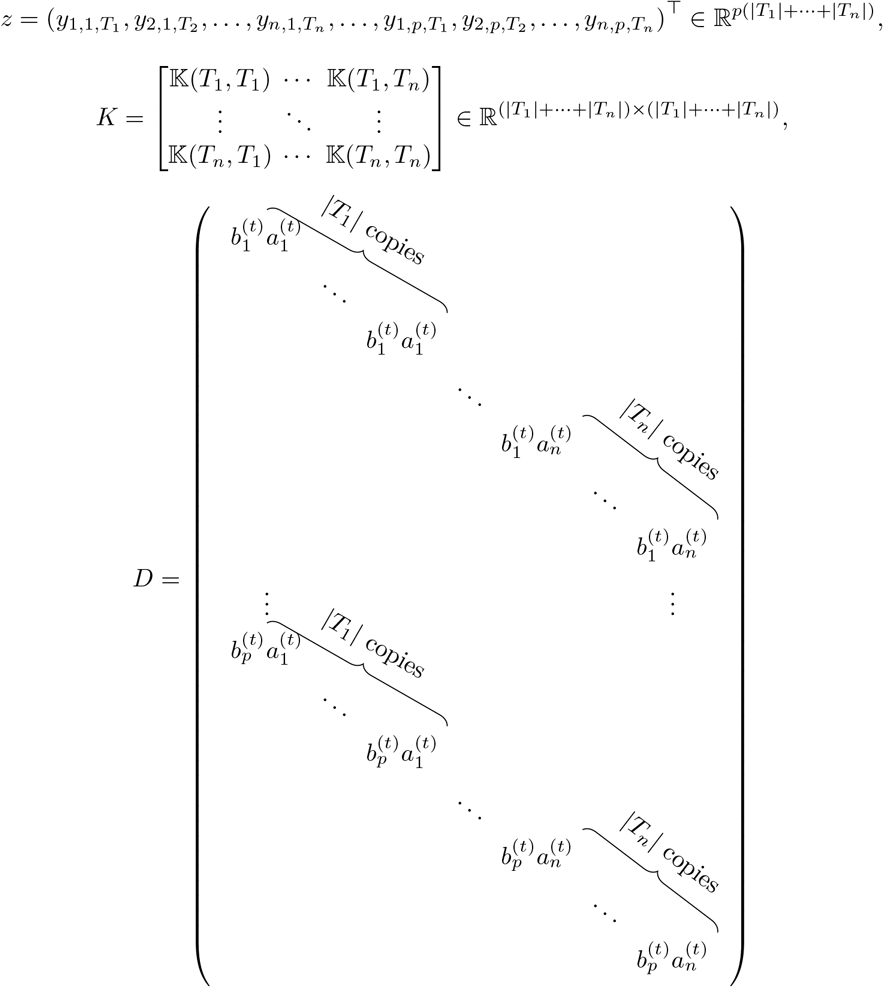

The solution to formula (9) is equivalent to

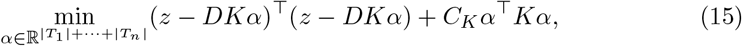

which can be solved by

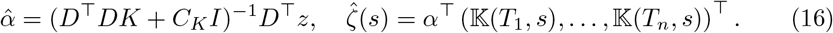

Here, *I* is the identity matrix, and *b*^(*t*)^ will cancel out in *D*^*T*^*D* because ||*b*^(*t*)^||_2_ = 1. *C*_*K*_ is a tuning parameter that makes *ζ*(*t*) smoother when *C*_*K*_ is larger. The implementation of TEMPTED allows users to choose the value of *C*_*K*_. We used *C*_*K*_ = 0.0001 for all our case studies and found it to work well for most datasets we analyzed with more than four time points. This is validated by a sensitivity analysis of *C*_*K*_ using the ECAM-based simulated data (Additional file 1: Fig. S2). Our results also indicate that while sample-level beta diversity derived from feature loadings is very insensitive to the choice of *C*_*K*_, a larger *C*_*K*_ is needed for a smaller number of time points to ensure good performance in the subject-level beta-diversity analysis.

### Tensor Reconstruction

In addition to estimating the low-rank components, we can also construct 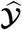 a low- rank approximation of the target tensor 𝒴, using the following method:

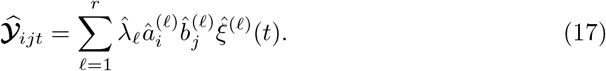

When mean subtraction is applied, the subtracted mean 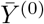 needs to be added back to 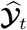 We evaluate the reconstruction accuracy of the decomposition in terms of normalized Frobenius norm:

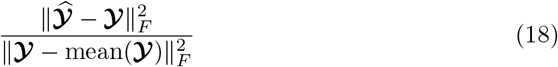

This reconstruction error ranges from 0 to 1 and decreases as the rank *r* increases for TEMPTED and FTSVD. It can guide the selection of *r* in the same fashion as (1 - percent of variance explained) in principal component analysis.

### Transfer of Dimension Reduction to New Data

Let 𝒴_train_ and 𝒴_test_ be two datasets consisting of the same features measured within the same time frame [0, *T*]. Suppose that TEMPTED decomposes 𝒴_train_ into *r* components, where the 𝓁th component has subject loading 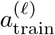, feature loading 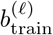, and temporal loading 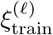. Assuming that 𝒴_train_ and 𝒴_test_ share the same feature loading and temporal loading, we can estimate the subject loading of 𝒴_test_ (i.e., 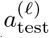) through one iteration of Step 2 of the TEMPTED algorithm, with 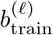 and 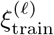 plugged in. The phenotype of the testing subjects will be predicted by applying classifiers to 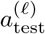 trained by 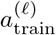. The prediction of phenotype for the testing data by such classifiers is purely out-of-sample since the dimension reduction is performed without any knowledge of 𝒴_test_, and classifiers trained by 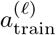 have no information from the testing data. The accuracy of such out-of-sample prediction is evaluated through data-driven simulation (Fig. 2 dashed lines).

### Case Study: Pax5 Mice Leukemia Data

The Pax5 dataset was published by [3] and deposited at https://qiita.ucsd.edu/study/description/11953, artifact ID 75878 [27]. The dataset we used here consists of fecal samples collected from 17 wild type and 27 Pax5 heterozygous mice that were treated with antibiotics at the beginning of the experiment, and 17 Pax5 mice developed leukemia during the study. Samples were subjected to 16S V4 short-read Illumina sequencing. Raw reads are deposited at ENA with accession PRJEB34720 [28]. Please refer to [3] for detailed sequence processing methods. Note that the original publication contains further amplicon sequence data and PacBio long read data not used here. Samples with *<* 35000 reads are removed. Mice with *<* 2 time points after filtering are removed. Time points were recorded as days from the end of antibiotic treatment and were divided by 7 to obtain weeks. TEMPTED uses ASVs appearing in *>* 5% of all samples and CTF uses ASVs appearing in *>* 10% of all samples. We used the top 3 components for all dimension reduction methods when analyzing this dataset. The smoothness parameter *C*_*K*_ was set to 10^−4^ for TEMPTED.

### Case Study: Shotgun Metagenomic Vaginal Data

The dataset used in this study was collected and published by [4] and deposited under dbGaP (study no. 20280; accession ID phs001523.v1.p1) [29]. It comprises a total of 705 vaginal samples obtained from 175 pregnant women visiting maternity clinics in Virginia and Washington. Among them, 40 women had preterm pregnancies, and the remaining 135 women had term pregnancies. To ensure data quality, subjects with less than two time points and MAGs that appeared in less than 5% of all the samples were removed from the analysis. For the time-specific analysis we used ALDEx2 with gestational weeks 5-36. A test was performed on weeks with at least 2 term and 2 preterm subjects. We conducted a rank-2 decomposition of this dataset using TEMPTED, focusing specifically on interpreting the first component. The smoothness parameter *C*_*K*_ was set to 10^−4^.

### Case Study: 16S Vaginal Data

The 16S data used in this study were published by [4, 5], and were deposited under BioProjects PRJNA393472 (subjects enrolled at the University of Alabama, Birmingham) and PRJNA821262 (subjects enrolled at Stanford University). Preprocessed data were obtained from [30]. We focus specifically on the second- and third-trimester pre-delivery vaginal samples from 141 term pregnancies and 49 pre-term pregnancies. Samples with *<* 40000 reads were removed, and ASVs (amplicon sequence variants) appearing in ≤ 5% of all the samples were removed as well. For time-specific analysis using ALDEx2, to ensure enough sample size at each time point, gestational days were floored to gestational weeks. No test was performed on Week 36 because it only contains three preterm subjects. We performed rank-2 decomposition of this dataset using TEMPTED, focusing specifically on interpreting the first component. The smoothness parameter *C*_*K*_ was set to 10^−4^.

### Data Driven Simulation

The ECAM dataset was published by [2] (Qiita ID 10249). Preprocessed data were obtained from [31]. Months 15 and 19 were removed due to their large amount of missingness. Operational taxonomic units (OTUs) appearing in ≤5% of the remaining samples were removed, as were samples with *<* 2000 reads. Subjects with fewer than nine time points were also removed, leaving 23 vaginally delivered infants and 17 Cesarean-delivered infants in the analysis. For *m* = 2, …, 9, *m* samples were randomly chosen from each subject to form the simulated dataset. Time points were recorded in days and used as is for TEMPTED. For CTF, microTensor, and TCAM, methods that demand input as a tabular tensor low-level or no missing time points, the order of the *m* samples was used as the time variable. Since CTF and microTensor can handle some missingness, we also ran them with time points rounded to month. The results are summarized in Additional file 1: Fig. S5, which shows worse performance for microTensor and slightly improved performance for CTF, but the superiority of TEMPTED remained obvious. For TEMPTED, the smoothness parameter *C*_*K*_ was set to 1, 0.1, 0.01, 0.005, for *m* = 2, 3, 4, 5 respectively, and 10^−4^ for *m* = 6, …, 9. These values of *C*_*K*_ match with the optimal *C*_*K*_ indicated by the sensitivity analysis in Additional file 1: Fig. S2.

The FARMM dataset was published by [16] (BioProject ID PRJNA675301). Preprocessed relative abundance data were obtained from [32]. Taxa appearing in fewer than 5 samples were removed. Samples with fewer than 5 taxa were also removed. Time point zero was removed because no subject in the vegan group has samples at time zero, making the missingness not at random. The remaining 15 time points were randomly dropped to achieve different percentages of missingness. Time points were used as is for TEMPTED, CTF, and microTensor, while TCAM cannot be run with such missingness. Simulated read counts are generated from multinomial distribution based on the observed relative abundance, which is equivalent to rarefication on the observed read counts. For TEMPTED, the smoothness parameter *C*_*K*_ was set to 10^−4^ in all settings.

We used the same central-log-transformed data as input for TEMPTED and TCAM, while CTF and microTensor take count data as input. The log-fold-change over baseline transformation used in the TCAM paper yields worse results for TCAM. The number of ranks *r* is set to 2 for all methods in the simulation analysis in Fig. 2.

### Software Usage

PERMANOVA was implemented using R package vegan. Logistic regression and random forest were implemented using R package stats and randomForest, respectively. AUC-ROC was calculated using R package PRROC. ALDEx2 was performed using R package ALDEx2 (1.32.0). CTF was performed using Python plugin gemelli (0.0.8).

## Declarations

## Availability of data and materials

The TEMPTED method is built into the R package “tempted” on CRAN (source codes and tutorial on GitHub https://github.com/pixushi/TEMPTED and Zenodo [33]) and the Python package “gemelli” (source codes and tutorial on GitHub https://github.com/biocore/gemelli and Zenodo [34]). All data analysis and simulation are performed using R and Python scripts through the R package reticulate. All processed data and codes used for this paper are available on GitHub https://github.com/pixushi/TEMPTED paper and Zenodo [35].

The Pax5 mice dataset [3] was deposited at Qiita with artifact ID 75878 [27] and in the ENA under accession PRJEB34720 [28].

The shotgun vaginal microbiome dataset [4, 20] was deposited under dbGaP (study no. 20280; accession ID phs001523.v1.p1) [29].

The 16S vaginal microbiome dataset [3, 5] was deposited under BioProjects PRJNA393472 (subjects enrolled at the University of Alabama, Birmingham) and PRJNA821262 (subjects enrolled at Stanford University). Preprocessed data were obtained from [30].

The ECAM dataset [2] was published under Qiita ID 10249. Preprocessed data were obtained from [31].

The FARMM dataset [16] was deposited under BioProject ID PRJNA675301. Preprocessed relative abundance data were obtained from [32].

## Authors’ contributions

PS, RH, and AZ conceived TEMPTED. PS designed and implemented TEMPTED in R. CM implemented TEMPTED in Python. PS, CM, and LS conceived the data-driven simulation scheme. PS performed the data-driven simulation and analyzed the 16S vaginal microbiome study. SJ and PS analyzed the Pax5 mice leukemia study. LS analyzed the shotgun vaginal microbiome study. PS, CM, and LS wrote the manuscript, with input from RK, SJ, KO, GB, and MS. RK supervised CM and LS. All authors reviewed and approved the final manuscript.

## Ethics approval and consent to participate

Not applicable.

## Consent for publication

Not applicable.

## Additional Files

Additional file 1: Supplementary Materials. This file provides additional details and results from numerical studies, including Figs. S1-S15 and Tables S1-S2.

Additional file 2: Taxonomy of top ASVs. This table contains the detailed taxonomic assignment of top ASVs identified by TEMPTED in the Pax5 Mice Leukemia Data and plotted in Figs. S14 and S15.

Additional file 3: ALDEx2 on 16S Vaginal Data. This table contains the result of ALDEx2 applied to the 16S vaginal data.

Additional file 4: ALDEx2 on Shotgun Metagenomic Vaginal Data. This table contains the result of ALDEx2 applied to the shotgun metagenomic vaginal data.

