## Supplementary figures and materials. for "TEMPTED: time-informed dimensionality reduction for longitudinal microbiome studies"

### Additional File 1

#### S1 Toy Example

We simulated a dataset with 100 features and three groups of subjects, each containing 20 subjects. For each subject, we randomly selected 20 time points from uniform distribution between 0 and 1. We considered three temporal trends: $f_1(t) = \frac{1}{2}\sin(2\pi t - \frac{1}{2}\pi) + \frac{1}{2}$, $f_2(t) = \text{atan}(10t - 5)/\pi + \text{atan}(5)/\pi$, and $f_3(t) = \frac{1}{2}\sin(4\pi t - \frac{1}{2}\pi) + \frac{1}{2}$. For Group 1, we simulated the absolute abundance of the first feature from $f_1(t)$ perturbed by a uniform distribution on (0, 0.5). For Groups 1 and 3, we simulated the absolute abundance of the second feature from $f_2(t)$ perturbed by a uniform distribution on (0, 0.25). For Groups 1, 2, and 3, we simulated the absolute abundance of the third feature from $f_3(t)$ perturbed by a uniform distribution on (0, 0.5). The absolute abundance of the remaining features was simulated from a uniform distribution on (0, 0.25). We then calculated the relative abundance and used them as proportion parameters in a multinomial distribution to generate counts. The sequencing depth was generated from an exponentially transformed normal distribution with a mean of 10 and a standard deviation of 0.5, resulting in minimum, median, and maximum depths of 5K, 22K, and 113K, respectively.

#### S2 Sensitivity analysis of model parameters

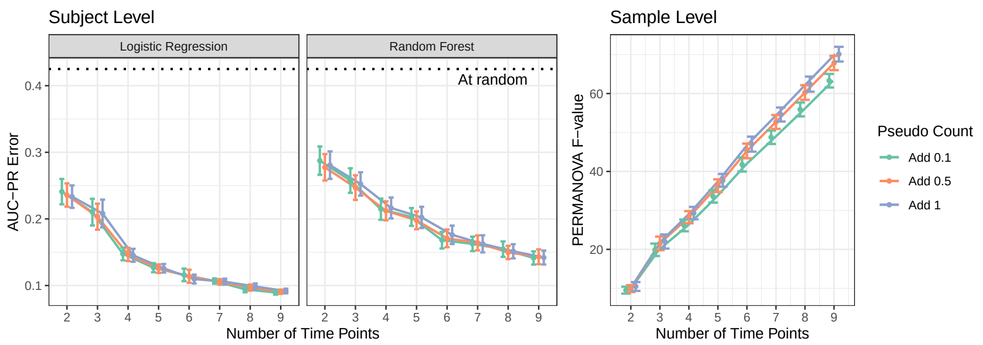


**Fig. S1** Impact of the choice of pseudocount used in centered log-ratio transformation. The simulation setting is the same as the ECAM-based simulation in Fig. 2(a-b). While using the pseudocount of 0.1 has slightly worse performance, choosing between 0.5 and 1 has little difference.

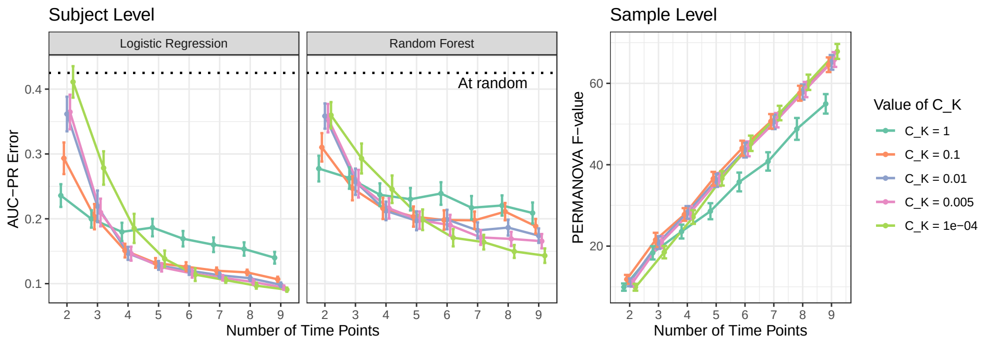


**Fig. S2** Impact of the choice of smoothness parameter $C_K$ used in TEMPTED. The simulation setting is the same as the ECAM-based simulation in Fig. 2(a-b). While sample-level beta diversity derived from feature loadings is very insensitive to the choice of $C_K$, a larger $C_K$ is needed for a smaller number of time points to ensure good performance in the subject-level beta-diversity analysis.

#### S3 Effect of Mean Subtraction

TEMPTED can be implemented with or without mean subtraction before the CP-type decomposition. In this section, we illustrate the effect of mean subtraction through ECAM-based and FARMM-based simulation under a set of parameter settings.

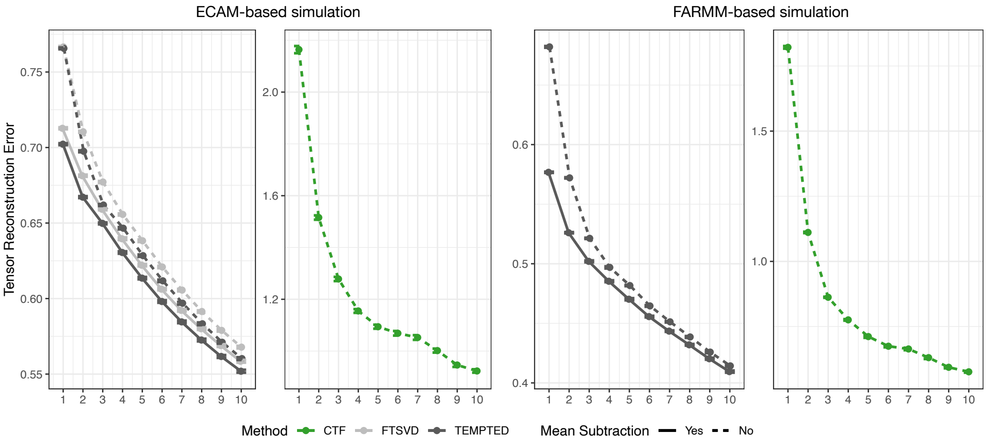


**Fig. S3** Comparison of reconstruction error between TEMPTED and other methods for different choices of rank parameter $r$. The simulation setting is the same as Fig. 2 with ECAM-based simulation at the number of time points equal to nine and FARMM-based simulation at the sequencing depth of 500K and percent of missingness being 20%. TCAM was not included in the benchmark because its decomposition retains a degree-of-freedom of $(n+p)*$(number of time points) for its first component. In contrast, TEMPTED, FTSVD, and CTF employ a CP-type decomposition that retains a degree-of-freedom of $n + p +$ (number of time points) for each component, which results in a significantly lower order of magnitude for the degree of freedom compared to TCAM. MicroTensor was excluded from the benchmark because it models counts directly with Dirichlet-Multinomial distribution and decomposes the underlying parameters instead of the observed data. Its reconstruction is focused on the proportion data. In contrast, TEMPTED, FTSVD, and CTF aim to minimize the $\ell_2$ loss for centered log-ratio transformed data and are not suitable for reconstructing proportions. Due to these differing reconstruction goals, comparing the accuracy of microTensor with the other methods is not meaningful.

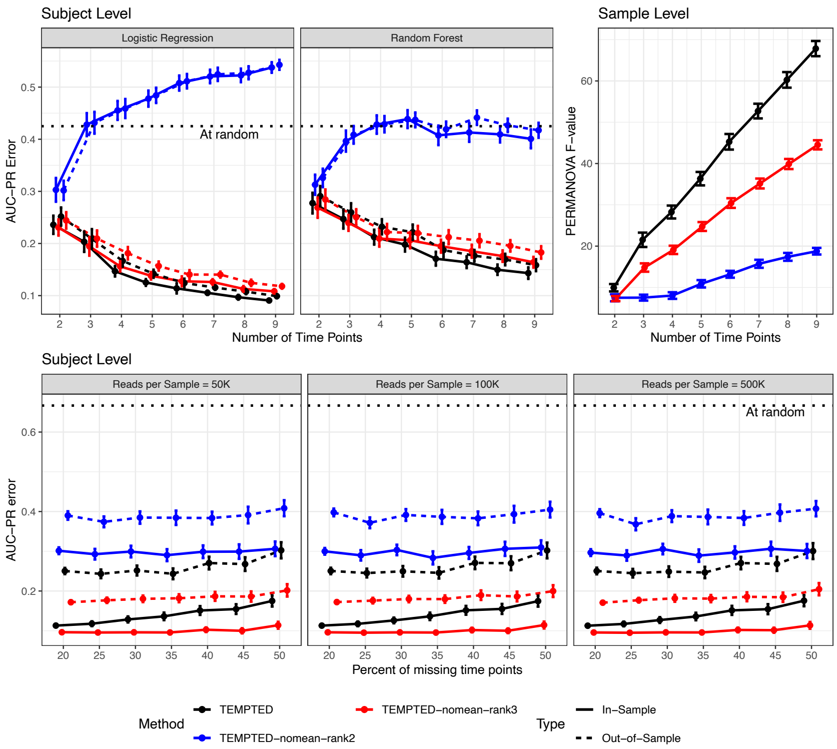


**Fig. S4** Comparison of group structure identification using TEMPTED with and without mean subtraction. The simulation setting is the same as Fig. 2. Black lines represent TEMPTED with mean subtraction at rank=2. Blue lines and red lines represent TEMPTED without mean subtraction at rank=2 and rank=3, respectively.

#### S4 ECAM-Based Simulation with Month as Time

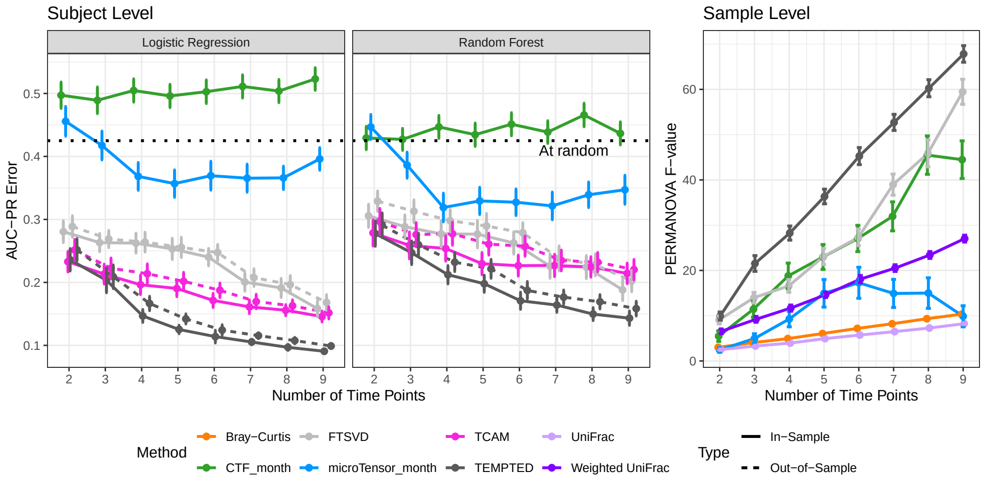


**Fig. S5** Recreation of Fig. 2 with microTensor and CTF run using month of age as time points.

#### S5 Temporal Sampling of Studies

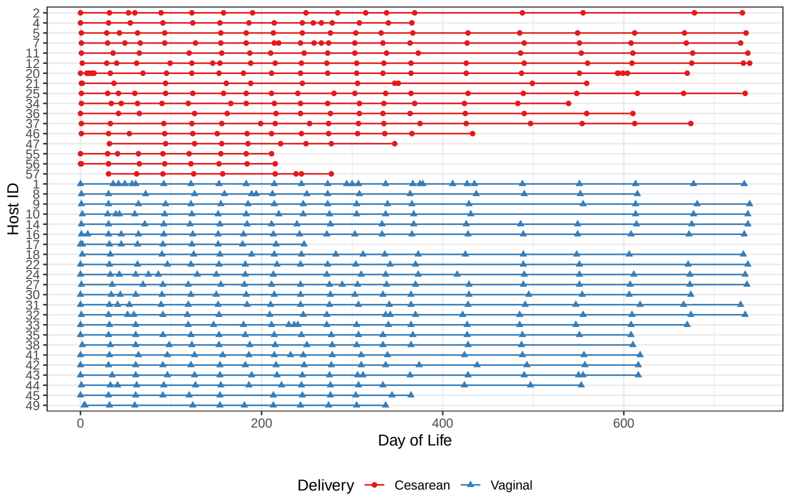


**Fig. S6** Temporal sampling of ECAM infant gut microbiome data. Each row represents one host, and each point represents one visit.

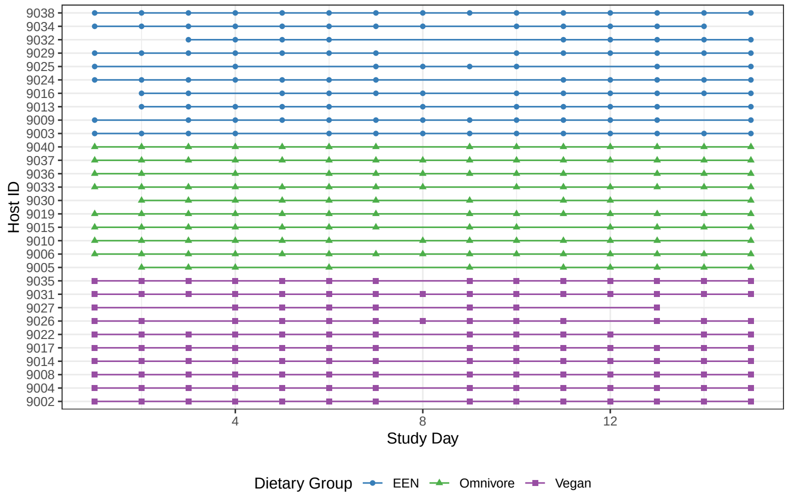


**Fig. S7** Temporal sampling of FARMM gut microbiome data

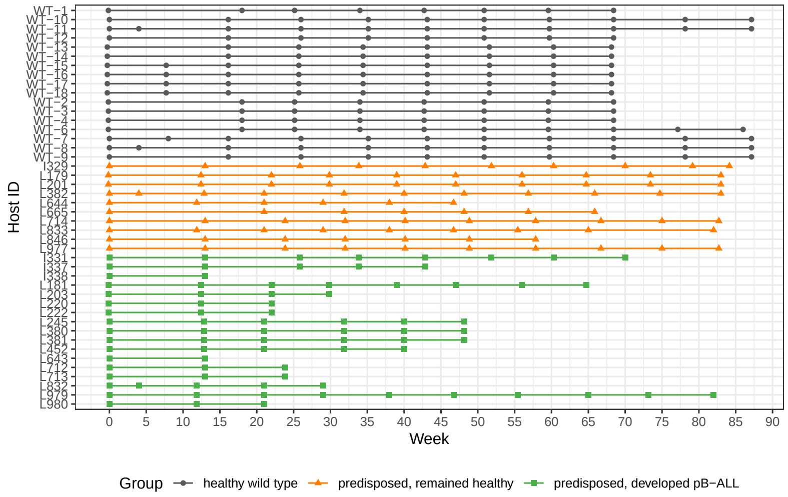


**Fig. S8** Temporal sampling of Pax5 mice leukemia data

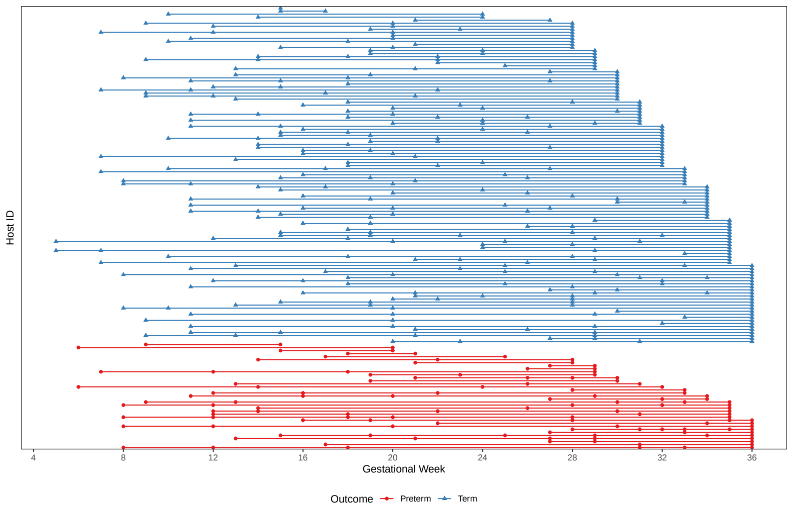


**Fig. S9** Temporal sampling of metagenomic vaginal microbiome data.

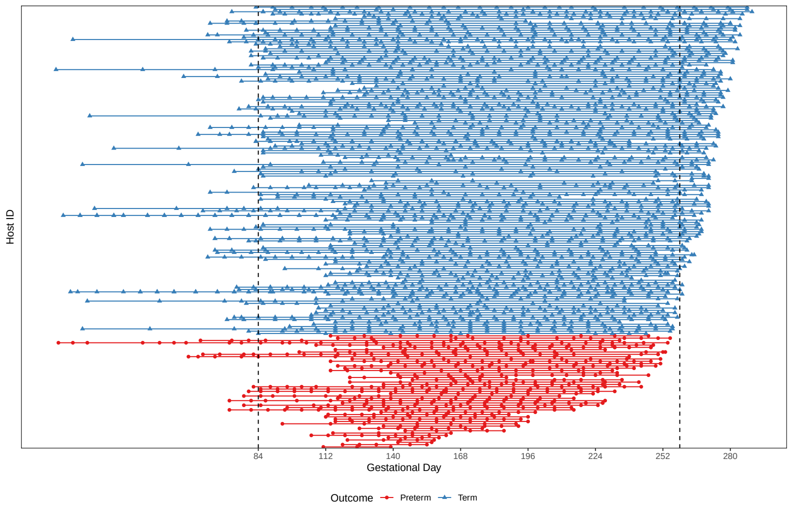


**Fig. S10** Temporal sampling of 16S vaginal microbiome data. Only second- and third-trimester samples, i.e. samples between the two dashed vertical lines, were used.

#### S6 Analysis of Pax5 Mice Leukemia Data

##### S6.1 Analysis with Other Methods

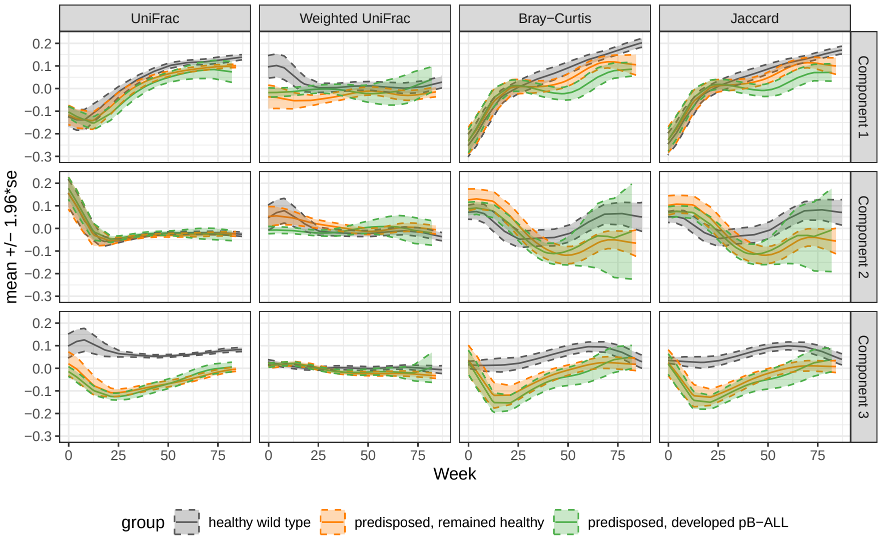


**Fig. S11** Trajectories of first three PCoA components for various distance metrics on the Pax5 mice leukemia data. Mean and error bands representing 1.96 standard errors of the mean are smoothed using kernel regression.

| | bray | jaccard | unifrac | wunifrac | TEMPTED | CTF |
|---|---|---|---|---|---|---|
| p-value | 0.143 | 0.143 | 0.342 | 0.798 | 0.001 | 0.032 |
| F-value | 1.996 | 1.930 | 0.956 | 0.145 | 6.922 | 3.707 |

**Table S1** PERMANOVA test between leukemia outcomes for 35-70 week samples among Pax5+/− mice

| Component | bray | jaccard | unifrac | wunifrac | TEMPTED | CTF |
|---|---|---|---|---|---|---|
| 1 | 0.007 | 0.006 | 0.135 | 0.672 | | 0.029 |
| 2 | 0.613 | 0.523 | 0.131 | 0.613 | 1.7e-5 | |
| 3 | 0.491 | 0.430 | 0.849 | 0.759 | | |

**Table S2** Wilcoxon test between leukemia outcomes for 35-70 week samples among Pax5+/− mice for each component

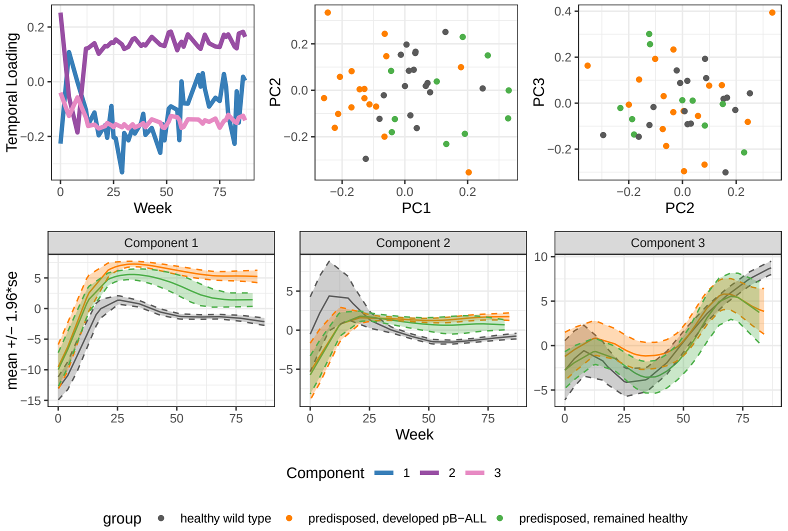


**Fig. S12** Results from CTF on the Pax5 mice leukemia data. The CTF results have an inconsistent separation of study groups between subject trajectories (predisposed pB-All > predisposed healthy > wildtype) and subject loadings (predisposed pB-All > wildtype > predisposed healthy), making it difficult to associate microbial features with the separation of study groups.

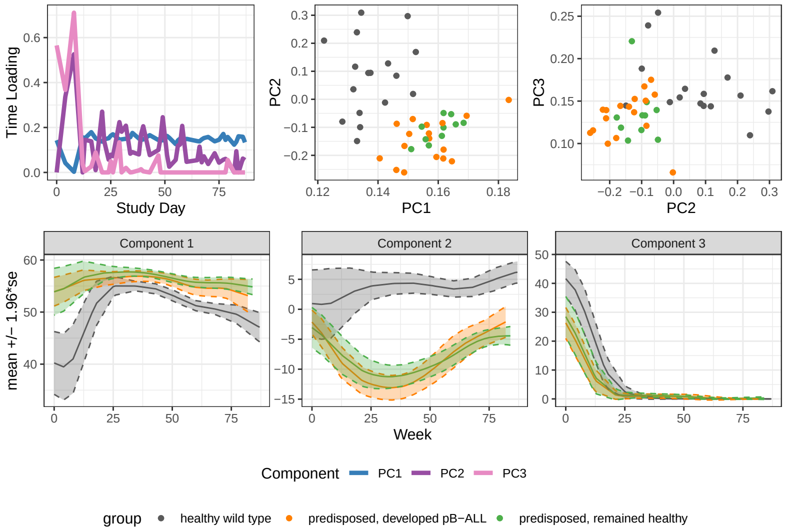


**Fig. S13** Results from microTensor on the Pax5 mice leukemia data. microTensor was unable to separate disease onset for predisposed mice in the first three components.

#### S6.2 Top ASVs identified by TEMPTED

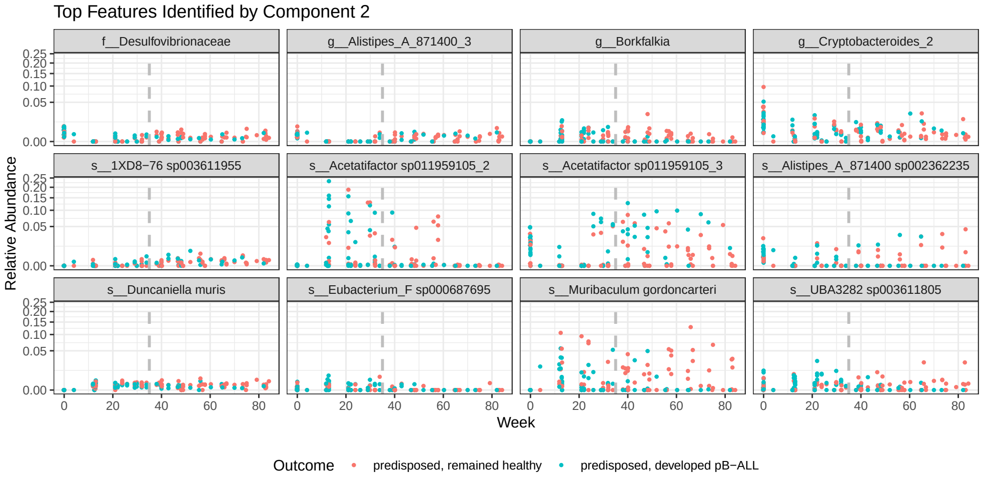


**Fig. S14** Relative abundance top ASVs identified by the second component of TEMPTED.

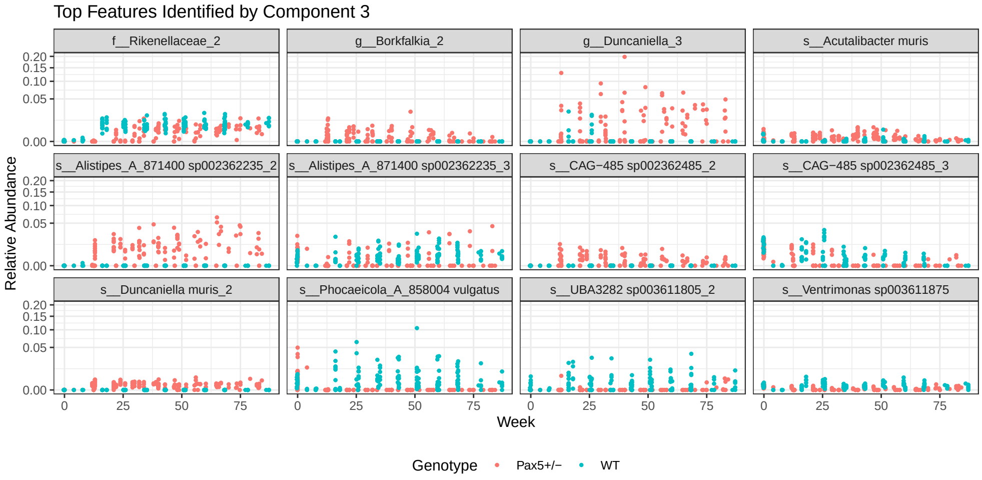


**Fig. S15** Relative abundance of top ASVs identified by the third component of TEMPTED.
